# Decoding Diabetes: Harnessing AI to Accurately Predict Real-Time and Future Blood Glucose Levels for Diabetes Management Using Diet, Exercise, Insulin Intake, and Heart Rate Variability

**DOI:** 10.1101/2025.11.02.686165

**Authors:** Riddhi Singhvi, Sanjay Singhvi

## Abstract

Continuous glucose monitoring (CGM) systems play a crucial role in diabetes care. Yet, they focus solely on blood glucose levels (BGL), neglect diet, exercise, and medication, and lack predictive capabilities, leaving patients and clinicians with reactive rather than proactive solutions. This study introduces Dual Temporal Recurrent Ensemble (DTRE), a novel AI model that bridges these gaps by enabling real-time BGL monitoring without a traditional CGM and forecasting BGL for 30 to 120 minutes into the future when integrated with CGM data. The model’s performance is based on two parallel branches of hybrid models that combine advanced machine learning architectures for accurate predictions and integrate key biomarkers like diet, exercise, insulin intake, heart rate (HR), and heart rate variability (HRV).

This pioneering study developed and validated DTRE on two diverse datasets, OHIOT1DM and D1NAMO, achieving exceptional forecasting accuracy: a Mean Absolute Relative Difference (MARD) of 7.6% at 30 minutes and 19.2% at 120 minutes, outperforming existing models by 13 to 41%. DTRE also surpassed commercial CGM systems for real-time BGL predictions by achieving a MARD of 7.17% on the high-frequency D1NAMO dataset compared to FreeStyle Libre 3 (7.9%) and Dexcom G7 (8.2%).

DTRE is the first AI-driven virtual BGM (vBGM) to integrate all four pillars of diabetes care and validate them on two diverse datasets. It offers a non-invasive, low-cost, and proactive solution. Its real-time insights can provide patients and clinicians with actionable data, transforming diabetes management for individuals who depend on insulin. Validation across diverse datasets underscores its potential for global application.

**Human Subject Research Statement:** This research did not utilize any human subjects. Pre-existing and published databases were used for this research.

## Introduction

In 2022, 8.75 million people worldwide had Type 1 diabetes (T1D) (IDF, 2022). Managing healthy blood glucose (BG) levels is a daily challenge for most patients. There are currently two primary methods for monitoring blood glucose (BG): continuous glucose monitoring (CGM) and fingerstick testing. Many struggle with the high cost of continuous glucose monitoring (CGM) systems or rely on invasive, infrequent fingerstick tests. Current tools do not provide comprehensive glucose management and low-cost real-time blood glucose (BG) prediction. This research utilizes Artificial Intelligence (AI) and machine learning (ML) to address these gaps and enhance the lives of millions of people worldwide.

The CGM systems have drastically improved glycemic control. They have become easier to use and more reliable. However, they have limitations because they cannot predict future BG levels. They also fail to address the other three pillars of diabetes care: diet, exercise, and medication (American Diabetes Association, 2024). This limits their ability to prevent BG fluctuations and optimize treatment.

Recent studies show that AI-driven models can predict glucose levels using physiological and behavioral data, offering an alternative to CGMs (Zhu et al., 2021). Kim et al. (2023) found that hybrid machine learning models combining these inputs improve prediction accuracy. However, challenges persist due to factors such as delays in food and insulin absorption, as well as a lag in interstitial tissue measurements (Guan, 2023).

Moreover, most current models rely heavily on CGM data and neglect non-invasive biomarkers such as heart rate (HR), heart rate variability (HRV), insulin dosages, and dietary patterns. Also, existing AI-based glucose prediction models primarily focus on short-term horizons of 30–60 minutes (Kim, 2020). A gap in long-term predictions (90–120 minutes) is critical for effective diabetes management (Smith et al., 2024).

HRV, which reflects autonomic nervous system activity, is an emerging biomarker for understanding glucose dynamics. The potential of HRV metrics in capturing blood glucose fluctuations could be an important, non-invasive tool in next-generation blood glucose monitoring technologies (Patel et al., 2023). This study integrates HRV metrics with diet, exercise, and insulin data for better glucose prediction. It is pioneering a holistic approach to glucose prediction that addresses the limitations of traditional CGM-based monitoring and models.

This research introduces a novel AI algorithm, the Dual Temporal Recurrent Ensemble (DTRE), that powers a Virtual Blood Glucose Monitor (vBGM). This vBGM operates in two modes: providing 30–120-minute predictions of future BG levels when CGM data is available, and offering real-time BG levels using non-invasive biomarkers without CGM. The algorithm leverages a dual hybrid AI architecture that integrates biomarkers such as diet, exercise, insulin, HR, and HRV to bridge gaps in existing models regarding the four pillars of diabetes management. Using preexisting, public domain datasets (OHIOT1DM and D1NAMO), the algorithm is rigorously trained and validated across both datasets. This research fills a critical void in current diabetes research. Additionally, the vBGM’s (Virtual Blood Glucose Monitor) ability to provide real-time predictions of BG without CGM offers a non-invasive, affordable alternative for those without CGMs

### Research Objective

This study aims to develop and validate a novel AI-driven Virtual Blood Glucose Monitor (vBGM) to enhance diabetes management. The vBGM provides real-time glucose predictions using non-invasive biomarkers, such as diet, exercise, insulin levels, and heart rate variability (HRV), thereby eliminating the need for CGM devices. Additionally, the vBGM provides future glucose forecasts at 30, 60, 90, and 120 minutes using CGM data, aiming to achieve superior accuracy compared to existing predictive models.

### Hypotheses

1. The novel AI algorithm is expected to achieve greater accuracy in predicting blood glucose levels at 30, 60, 90, and 120 minutes compared to existing models, particularly at the longer horizons of 90 and 120 minutes.
2. The vBGM will provide clinically relevant, real-time glucose predictions using non-invasive biomarkers, albeit with less accuracy than CGM-based predictions.
3. The D1NAMO dataset, with a higher data-collection frequency, will yield more accurate results than the OHIOT1DM dataset for real-time vBGM predictions.
4. The dual capability of the vBGM, with real-time and extended future predictions, will demonstrate clinical reliability in accordance with Clarke Error Grid Zone A/B standards, addressing critical gaps in current diabetes management tools.

## Materials and Methods

### Datasets

#### D1NAMO Dataset

The D1NAMO (Diabetes Monitoring) dataset was developed to support non-invasive glucose management research, collected at Hôpital Riviera-Chablais in Vevey, Switzerland (Dubosson, 2018), with ethical approval (CER-VD 53/15). It includes data from nine Type 1 diabetes (T1D) patients (six males and three females, aged 20–79) who wore the Zephyr BioHarness 3 for four days, capturing ECG signals at 250 Hz, breathing data at 18 Hz, and accelerometer readings at 100 Hz. Continuous glucose monitoring (CGM) data were recorded every five minutes using the iPro2 system, alongside annotations of 115 meals and 106 images for caloric and macronutrient content. The dataset provides approximately 450 hours of physiological recordings and 8,414 glucose measurements. Key challenges included the short monitoring duration, noisy data, and a small sample size. The dataset was split into 80% for training and 20% for testing for model development and validation.

#### OHIOT1DM Dataset

The OHIOT1DM (OHIO) dataset comprises long-term glucose data collected under real-world conditions over eight weeks at Ohio University (Marling, 2020). It includes 12 participants (seven males, five females, aged 20–80), categorized into three age groups: young adults (20–40 years), middle-aged adults (40–60 years), and older adults (60–80 years). The dataset comprises CGM readings every five minutes, insulin logs detailing basal and bolus doses, heart rate (HR) data, and self-reported information on meal timing, carbohydrate intake, exercise intensity, sleep quality, and stress events. It encompasses 115,000 glucose readings and 1,500 life events. Challenges included data noise, imbalance, and the synchronization of diverse streams, which were addressed through careful preprocessing and the use of predefined training and testing subsets.

#### Computational Environment

Based on Python 3.10.9, the CPython interpreter and GCC 11.2.0 are the compilers. The models were developed and evaluated on a Linux platform (kernel version: 6.8.0-49-generic-x86_64) with glibc 2.35. Key libraries included Keras 2.10.0 for building deep learning models, TensorFlow 2.10.0 as the backend, and scikit-learn 1.3.2 for preprocessing and machine learning algorithms. NumPy 1.24.4 facilitated numerical computations, and Pandas 1.5.3 handled data manipulation and visualization tasks using Matplotlib 3.7.0.

### Feature Engineering

Feature engineering was critical for transforming raw physiological data into meaningful inputs. Essential features included carbohydrates, insulin, heart rate variability (HRV) metrics, and circadian representations. Sine and cosine transformations of time data captured daily glucose variations and circadian rhythms, enabling the models to distinguish between morning and evening patterns.

HRV metrics, extracted from ECG signals using the PyHRV toolkit (Gomes, 2019), provided insights into autonomic nervous system activity. Recent advancements show the importance of feature engineering in developing AI-driven diabetes care solutions (Liu, 2024). The inclusion of metrics such as Root Mean Square of Successive Differences (RMSSD), the standard deviation of NN intervals (SDNN), Triangular Interpolation of NN interval histogram (TINN), and HRV TriIndex ensures that the models leverage rich physiological data for robust glucose prediction.

### Carbohydrate Absorption Dynamics Using a Piecewise Approach

The piecewise function used to model carbohydrate absorption is based on established computational models (El-Khatib et al., 2010) that simulate the time-dependent effects of dietary intake on glucose dynamics. This equation models the dynamics of effective carbohydrate absorption over time after a meal, divided into three phases. No carbohydrates are absorbed in the initial phase (0 ≤ t_meal_ < 2), where t_meal_ is the meal time. During the rising phase (2 ≤ t_meal_ < 12), the effective carbohydrate level C_cp_(t) increases, peaking at t_peak_, driven by the increasing rate of the curve (α_inc_) and total carbohydrates (C_meal_). In the declining phase (12 ≤ t_meal_ <48), C_cp_(t) decreases following a decay curve defined by the decreasing rate of the curve (α_dec_). This equation provides a structured representation of the dynamics of carbohydrate absorption and metabolism.

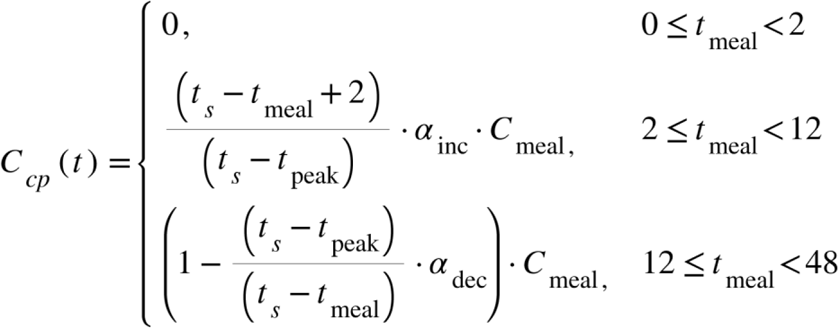

where:

*t*_*s*_ is the sampling time.
*t*_*meal*_ is the time when the meal is encountered.
*C*_*op*_ is the effective carbohydrates at any given time.
*C*_*meal*_ is the total amount of carbohydrates taken in a meal.
*t*_*peak*_ is the time when *C*_*op*_ reaches its maximum value → *C*_*op*_ ≈ *C*_*meal*_
*⍺*_*inc*_ is the increasing rate of the curve.
*⍺*_*dec*_ is the decreasing rate of the curve.

#### Active Insulin Dynamics

Insulin activity was modeled using exponential decay for rapid-acting (4 hours) and long-acting (24 hours) insulin, creating an “Insulin on Board” (IOB) feature as an input to the model. The insulin rate (*R*_*insulin*_(*t*)) is the combined effect of doses distributed over their respective action times: fast-acting insulin (*R*_*fast*_(*t*)) was distributed over 4 hours (*T*_*fast*_ = 60 ⋅ 60 ⋅ 4), and Slow-acting insulin (*R*_*slow*_(*t*)) was distributed over 24 hours (*T*_*slow*_ = 60 ⋅ 60 ⋅ 24). Contributions from overlapping doses were summed at each time step, ensuring an accurate representation of insulin dynamics.

Mathematical Representation:

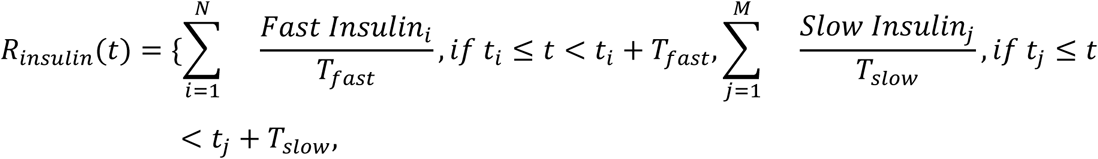

Where:

*t*_*i*_: Time of the *i*-the fast insulin dose.
*t*_*j*_: Time of the *j*-th slow insulin dose.
*T*_*fast*_, *T*_*slow*_: Duration over which each dose is distributed.

### Canonical Correlation Analysis

Canonical Correlation Analysis (CCA) was employed to examine the relationship between physiological features and glucose levels over a 180-minute interval in all patients. Glucose level was the dependent variable, and the independent variables varied according to the dataset. The preprocessing ensured the data was ready for the CCA model, enabling it to identify relationships between physiological features and glucose levels. The analysis identified linear combinations of features that maximized correlation with glucose changes. Variables were normalized, and outliers were removed. Python was used for analysis, and canonical variates were tested for significance.

### Model Description: Dual Temporal Recurrent Ensemble (DTRE)

This research presents an innovative hybrid modeling framework, the Dual Temporal Recurrent Ensemble (DTRE), designed to accurately forecast blood glucose levels using multiple biomarkers. The DTRE framework integrates two parallel hybrid branches: a virtual Blood Glucose Model and a Forecasting Model branch.

### Virtual Blood Glucose Model (vBGM) Branch

The first model, the Virtual Blood Glucose Model (VCGM), adopts a hybrid GRU-XGBoost approach. The GRU component captures long-term trends in glucose dynamics using two layers with 128 and 64 units, respectively, and incorporates 20% dropout for regularization. Residuals from the GRU are refined using XGBoost, configured with 100 trees, a maximum depth of 10, and a learning rate of 0.1. This hybrid model approach utilizes the GRU’s strength in identifying trends and the XGBoost’s capacity to capture non-linear variations, thus delivering strong performance across various temporal patterns.

### Forecasting Model Branch

The second model, an ensemble LSTM-GRU, uses the strengths of both model architectures by processing parallel branches. LSTM and GRU layers were fine-tuned using Keras Tuner, with units ranging from 64 to 512, dropout rates between 0.1 and 0.5, and L2 regularization parameters ranging from 1 × 10^−5^ to 1 × 10^−3^. The outputs of these branches are concatenated and passed through a dense layer to generate future glucose predictions. Model optimization was achieved using the Adam optimizer with a learning rate tuned between 1 × 10^−4^ and 1 × 10^−2^, while the Huber loss function was employed to minimize RMSE.

### Model Integration and Training

Dataset Preprocessing: Preprocessing included normalization, handling missing data, and aligning temporal data streams for the D1NAMO and OHIO datasets. Bhimireddy et al. (2020) describe similar preprocessing approaches, including feature alignment and handling imbalanced data streams, as critical for ensuring the robustness of glucose prediction models. Marling and Bunescu (2020) detailed the preprocessing techniques applied to the datasets, such as interpolating missing CGM data points and aligning self-reported information with CGM readings.

Additionally, noise reduction for high-frequency physiological data in the D1NAMO dataset followed the guidelines of Butt, Khosa, and Iftikhar (2023), which suggest using filtering techniques to extract features from ECG and accelerometer signals reliably.

Extensive data preprocessing was performed to ensure robust model performance. Features were scaled using MinMax normalization, and sequence preparation involved a sliding window approach with a window size of six. Rolling-window cross-validation with a 10-fold TimeSeriesSplit was applied to maintain temporal consistency, training on earlier segments and validating on later ones. This approach ensured the models were rigorously evaluated across diverse temporal patterns while minimizing the risk of overfitting. Hyperparameter optimization was conducted using Keras Tuner’s Hyperband search algorithm, enabling the identification of the optimal configurations for each architecture.

The DTRE framework’s advanced, novel hybrid structure and optimization show strong promise for predicting current and future blood glucose levels. Its incorporation of trend analysis, non-linear error adjustment, and time-based validation establishes a new method for predicting blood glucose accuracy in diabetes management.

### Evaluation Metrics

The machine learning models were evaluated using the following standard metrics:

1. Root Mean Squared Error (RMSE): RMSE quantifies the average magnitude of errors between predicted and actual glucose values, placing greater weight on larger deviations. It is a commonly used metric for evaluating the accuracy and reliability of predictive models across various domains, including glucose monitoring (Willmott & Matsuura, 2005).
2. Mean Absolute Relative Difference (MARD): MARD calculates the average relative error between predicted and actual glucose values, expressed as a percentage. It is considered a clinically relevant metric for evaluating the accuracy of blood glucose predictions (Freckmann et al., 2012).
3. Clarke Error Grid Analysis: This metric evaluates the clinical relevance of glucose predictions by categorizing errors into zones (A to E). Predictions in Zone A are considered clinically accurate, while in Zones C to E, errors may lead to wrong treatment decisions, highlighting the importance of predictive safety and reliability (Clarke et al., 1987).

### Ablation Analysis

A reverse ablation analysis of the forecasting and vBGM models was performed to evaluate the contribution of individual components by systematically removing them and measuring their impact on performance using MARD and RMSE as evaluation metrics. This analysis helped identify critical factors driving the model’s accuracy and provided a deeper understanding of the model’s parameters.

## Results

### Canonical Correlation Analysis (CCA) Analysis for D1NAMO Data

Canonical Correlation Analysis (CCA) revealed significant relationships between physiological metrics and glucose levels over 180 minutes. The main findings are summarized in Table 1 below:

**Table 1.** CCA analysis for OHIO dataset.

| Variable | Mean | Std Deviation |
| --- | --- | --- |
| HR | -6.09 | 15.48 |
| Physical | 0.02 | 0.34 |
| Operative Carbs | 0.14 | 0.12 |
| Insulin | 0.07 | 0.25 |
| Sin_Time | -0.13 | 0.23 |
| Cos_Time | -0.10 | 0.23 |
| Skin Temp. | -5.87 | 14.79 |
| Sleep | 0.17 | 0.12 |
| Steps | -0.16 | 0.92 |

The variables were normalized to ensure consistency, with most having means near zero. Operative Carbs, Insulin, Sleep, and Physical Activity showed low variability, indicating stable patterns across the dataset. Heart Rate (HR), Skin Temperature, and Steps showed greater variability, reflecting their dynamic nature and their ability to capture glucose changes. Sin_Time and Cos_Time accounted for daily glucose patterns. High-variability features were more influential in capturing glucose fluctuations, while low-variability features reflected consistent baseline effects.

Table 2 summarizes the Canonical Correlation Analysis (CCA) for the D1NAMO dataset, showing the relationship between biomarkers and glucose levels. Heart Rate (HR) and Breathing Rate (BR) show moderate positive correlations, with Patient 004 consistently exhibiting the highest values for both metrics. Other metrics, such as SDNN and NN_50, show weak average correlations but higher variability, with Patient 007 often showing the strongest correlations.

**Table 2.** CCA analysis for the D1NAMO dataset.

| Metric | Avg. correlation | Std deviation | Highest Correlation (Patient ID) |
| --- | --- | --- | --- |
| HR | 0.1173 | 0.0454 | 0.1993 (Patient 004) |
| BR | 0.0732 | 0.1053 | 0.1787 (Patient 004 ) |
| Sdnn | 0.0086 | 0.2125 | 0.179 (Patient 007) |
| Rmssd | -0.2513 | 0.3913 | 0.1228 (Patient 007) |
| NN_50 | 0.0097 | 0.2371 | 0.1843 (Patient 004) |
| pNN_50 | 0.0189 | 0.1603 | 0.1214 (Patient 004) |
| MeanHR | -0.0690 | 0.2347 | 0.2121 (Patient 007) |
| MeanRR | 0.0102 | 0.1936 | 0.2379 (Patient 004) |
| SDHR | -0.1071 | 0.3478 | 0.2 (Patient 007) |
| HRVTriIndex | 0.0037 | 0.1091 | 0.0978 (Patient 002) |
| TINN | 0.0114 | 0.1135 | 0.1557 (Patient 004) |
| R_Peaks | -0.2871 | 0.3309 | 0.0758 (Patient 002) |
| Insulin Fast | -0.8427 | 2.2135 | 0.092 (Patient 001) |
| Insulin Rate | 0.1240 | 0.0869 | 0.2856 (Patient 007) |
| Activity Level | 0.0638 | 0.1389 | 0.2225 (Patient 007) |
| Calories Exp | -0.0499 | 0.2076 | 0.0838 (Patient 006) |

HR and BR consistently show moderate positive correlations, with Patient 004 exhibiting the highest values for these metrics. The insulin rate also demonstrates strong positive correlations, validating its importance as a key predictor of glucose levels.

In contrast, RMSSD and R_Peaks generally show weak or negative average correlations, indicating limited predictive value. However, RMSSD shows a positive correlation for Patient 007. Insulin Fast shows a strong negative correlation, suggesting its limited utility in glucose prediction.

Patient-specific variations are evident, with Patients 004 and 007 showing the strongest correlations across several metrics. These differences highlight the individualized nature of glucose dynamics.

Activity Level and Calories Expended show small overall correlations, but Patient 007 shows the greatest impact on Activity Level. Meanwhile, MeanHR shows a negative average correlation, whereas MeanRR shows a small positive correlation, further complicating the interplay of factors influencing glucose levels.

### Forecasting Model Performance

The performance of the DTRE model was compared with several well-established models, including those developed by Sun et al. (2018), Gülesir et al. (2018), and Idriss et al. (2019). These previously published models represent the leading methodologies for glucose prediction, utilizing recurrent neural networks (RNNs) in conjunction with traditional machine learning approaches. These baseline models were selected because they were created using the OHIO dataset and focus on predicting glucose levels over different time horizons, thus aligning well with the objectives of this research. Importantly, these models predominantly use CGM data and do not fully incorporate non-invasive biomarkers, highlighting a significant gap that the DTRE model aims to address. To the best of our knowledge, no long-term forecasting model based on the D1NAMO dataset has been attempted in the published literature.

### Performance Comparison

Table 3 compares the proposed DTRE model with several published models across multiple glucose prediction horizons (30, 60, 90, and 120 minutes) using CGM data. Performance was evaluated using key metrics: MARD, RMSE, and the percentage of predictions in Zone A of the Clarke Error Grid.

**Table 3.**
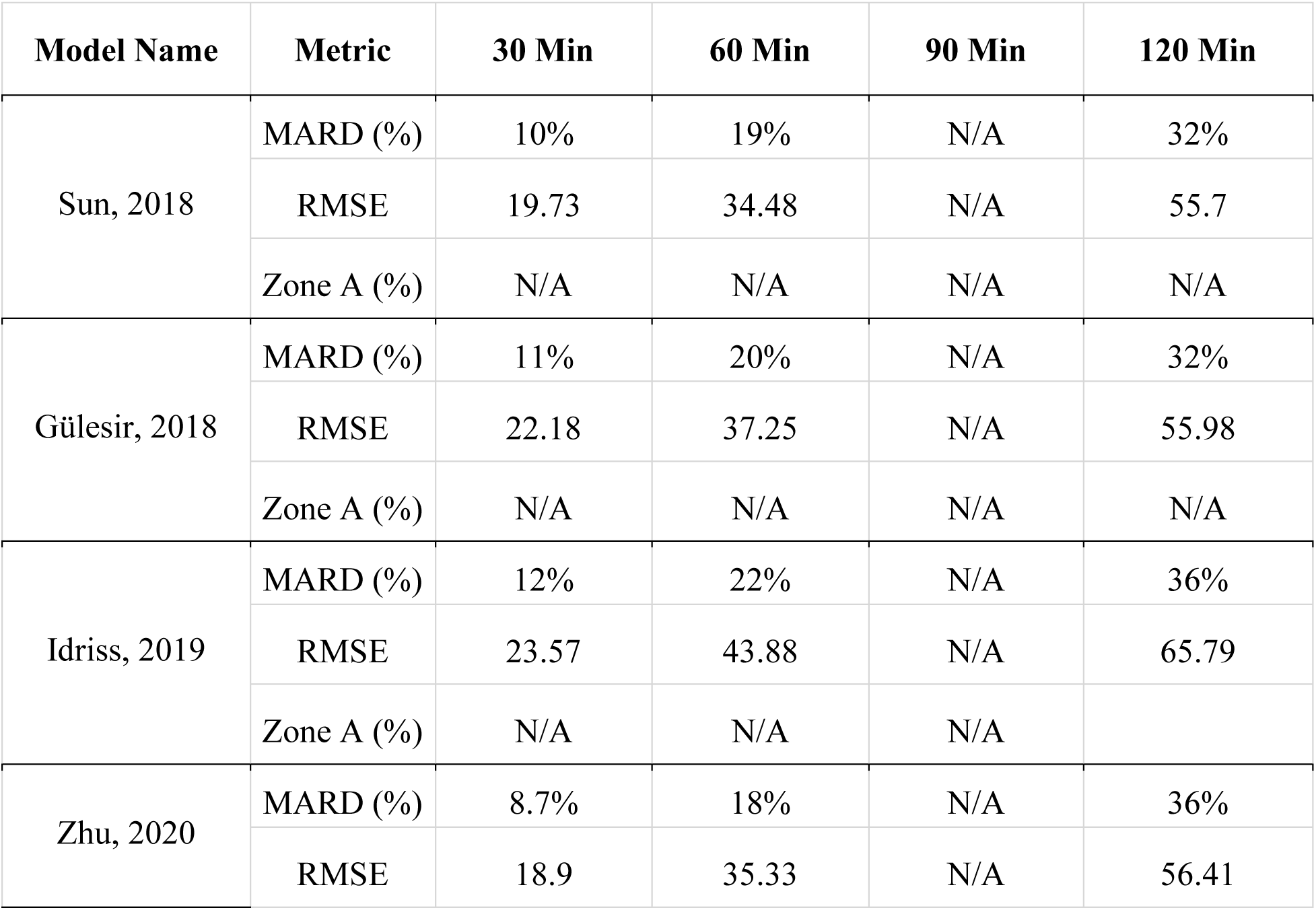
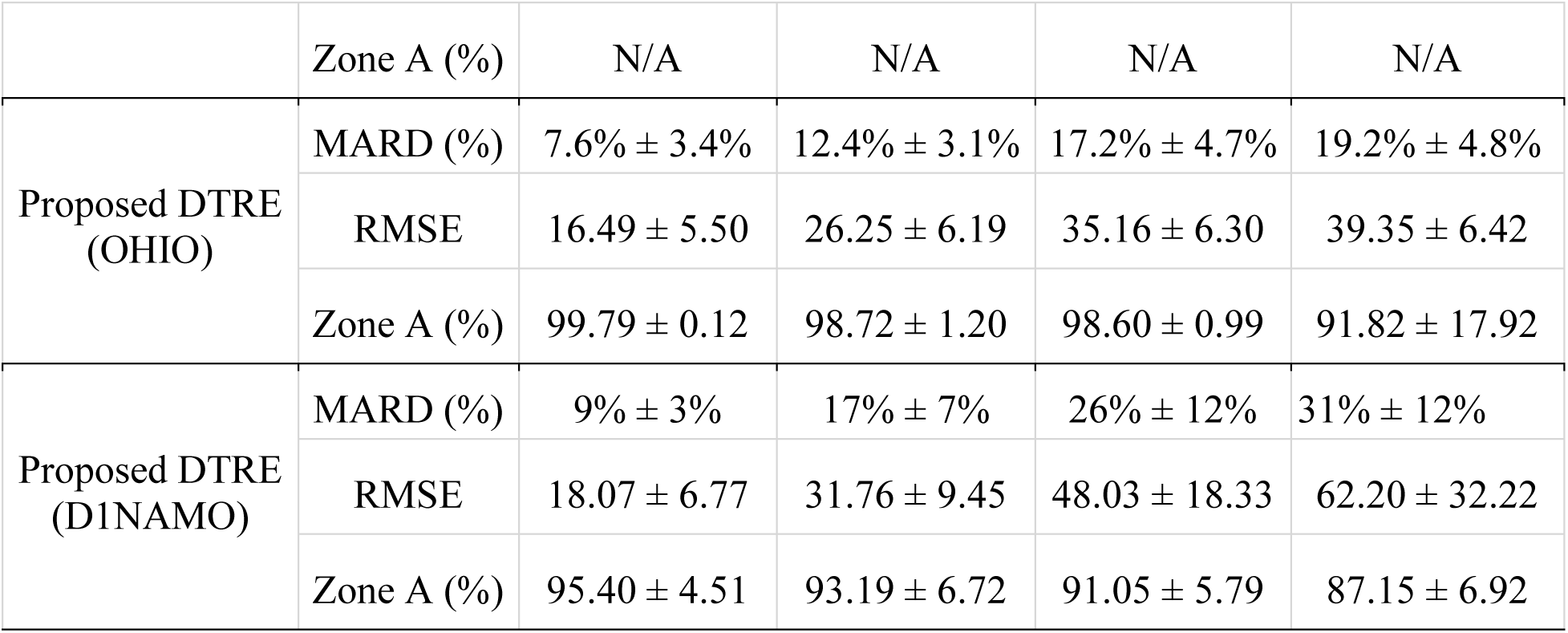
Comparison of different published models with Dual Temporal Recurrent Ensemble.

The DTRE model significantly outperforms all published models across all prediction horizons for MARD, achieving the lowest error rates. For instance, at the 30-minute horizon, the DTRE model achieves a MARD of 7.6% ± 3.4%, compared to 8.7% by Zhu’s model. Similarly, the DTRE consistently delivers lower RMSE values, reflecting improved accuracy in glucose level predictions. For Clarke Error Grid Zone A performance, the DTRE model achieves 99.79% accuracy in Zone A at the 30-minute horizon and 91.82% accuracy even at 120 minutes, indicating high clinical reliability.

Figure 1 illustrates the model’s ability to accurately predict blood sugar levels for Patient 570 over 90 minutes, demonstrating its strong performance over time. Predictions for other time intervals (30, 60, and 120 minutes) are included in the Appendix.

**Figure 1.**
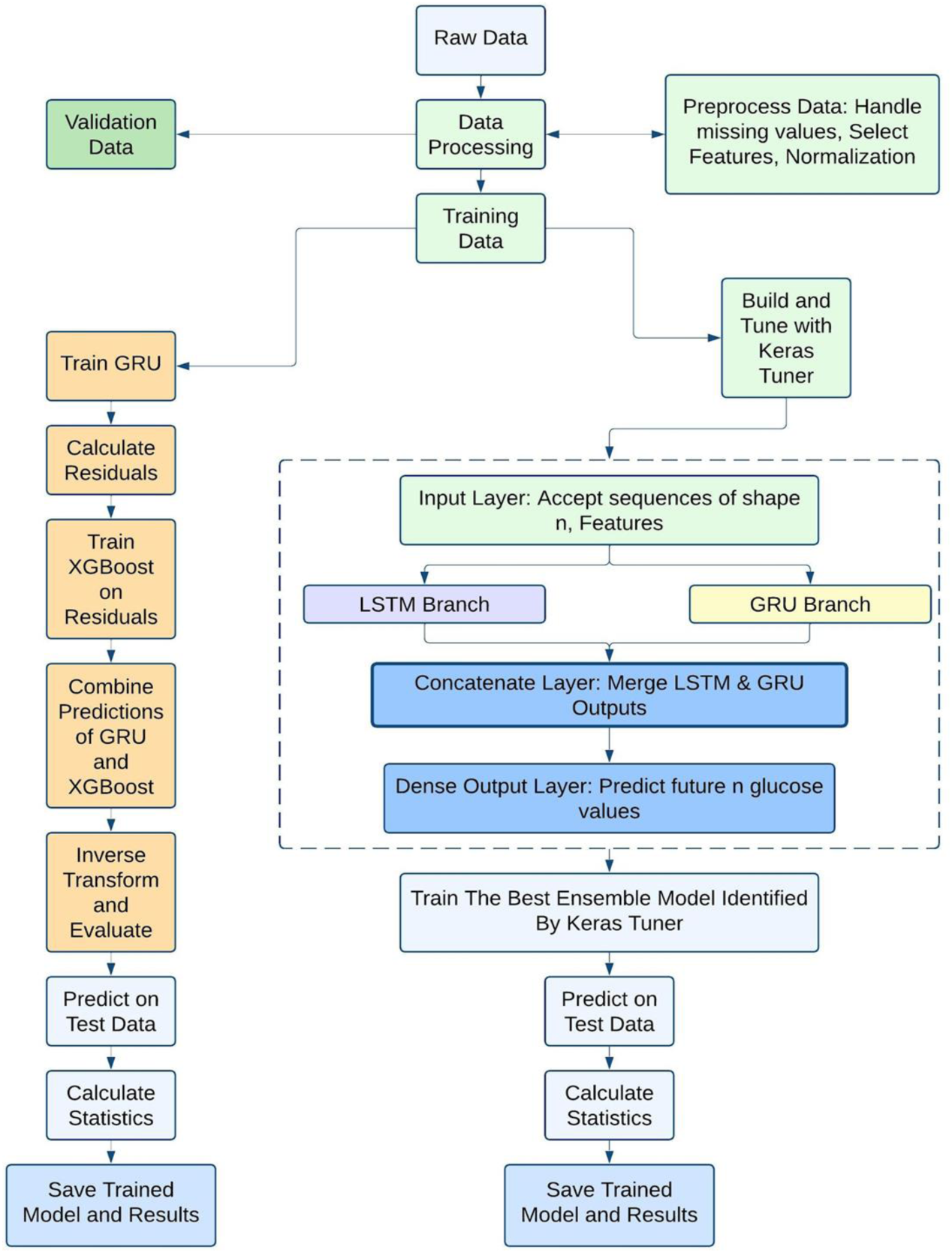
Flowchart of DTRE data processing.

**Figure 2.**
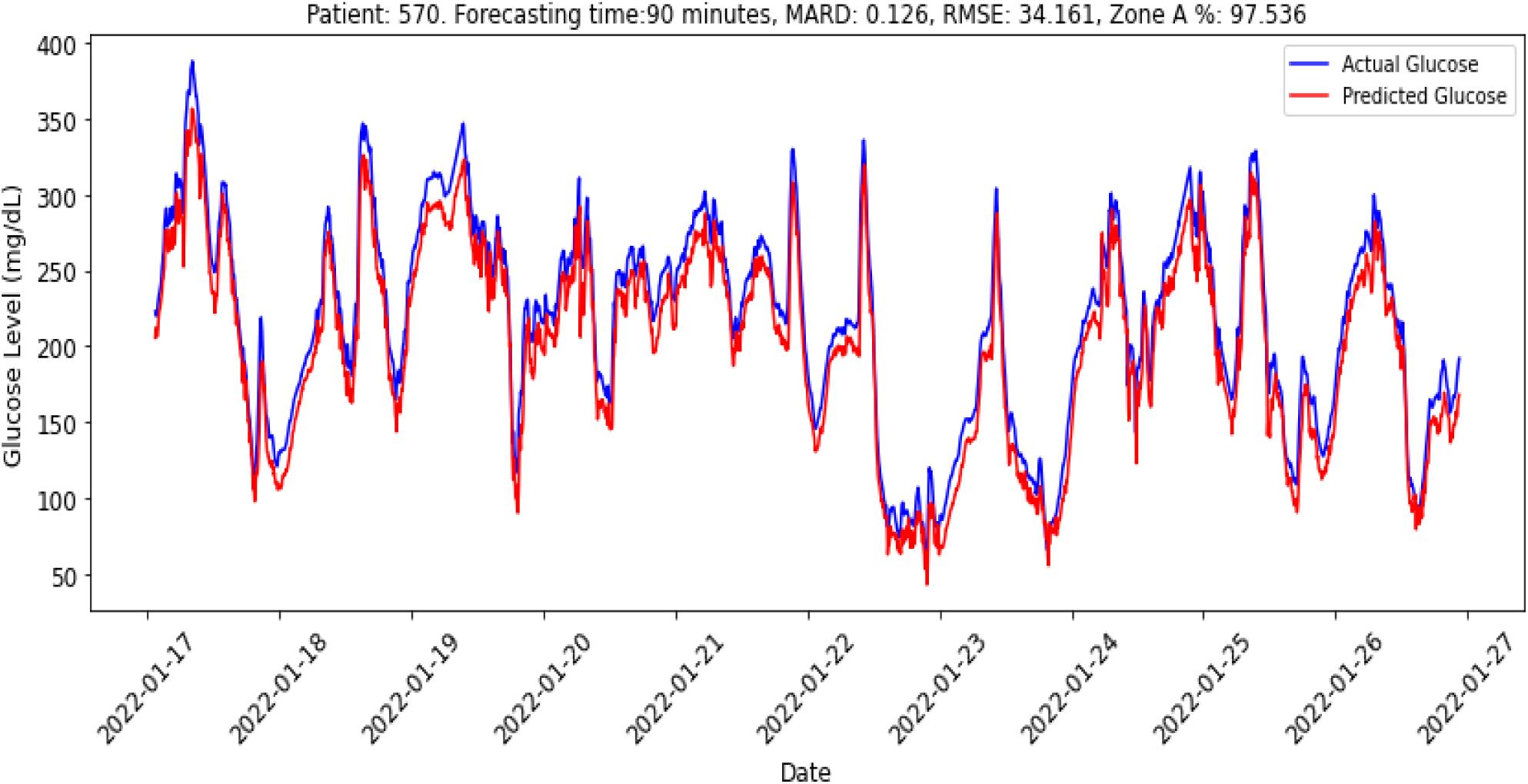
Performance of the DTRE model for 90-minute future forecasting for the OHIO dataset for Patient 570.

Figure 3 shows the 90-minute prediction results for Patient 006 using the D1NAMO dataset. The predicted glucose levels closely match the actual levels, showing the model’s accuracy. It successfully tracks the ups and downs, including peaks and dips, demonstrating its ability to handle changing glucose patterns over time.

**Figure 3.**
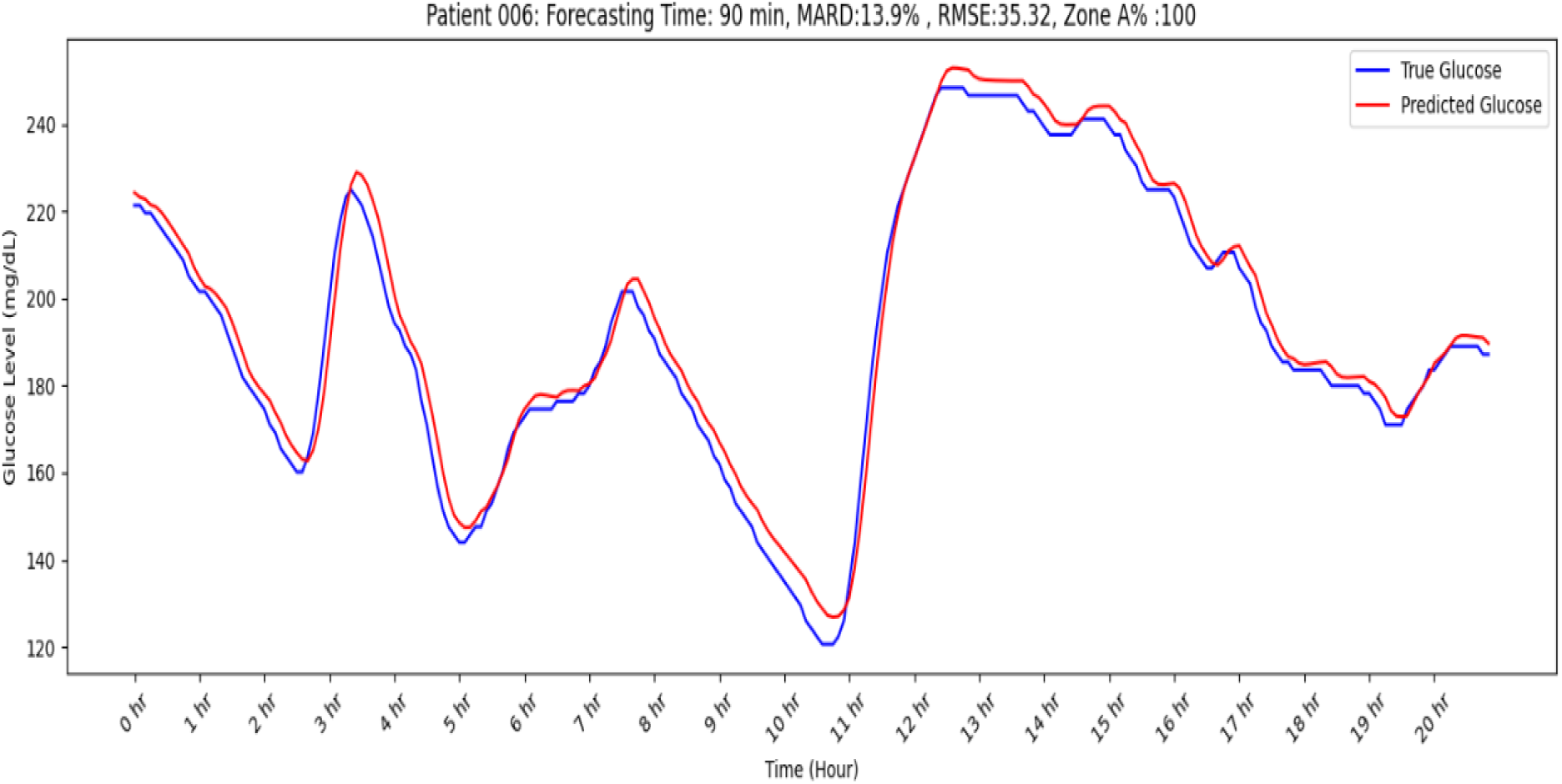
Performance of the DTRE model for 90-minute future forecasting. D1NAMO dataset for Patient 006.

### Virtual Blood Glucose Monitor (vBGM) Real-time Performance

Table 4 compares the DTRE model’s ability to predict blood sugar in real time across two datasets, without CGM. The model relied on past training and recent biomarkers for its predictions.

**Table 4.** Comparison of performance of vBGM on different datasets.

| Dataset | Patient ID | MARD | RMSE | Zone A | Zone B | Zone C | Zone D | Zone E |
| --- | --- | --- | --- | --- | --- | --- | --- | --- |
| OHIO | 563 | 16.92 | 34.71 | 69.60 | 28.75 | 0.18 | 1.47 | 0 |
|  | 570 | 22.67 | 56.93 | 53.47 | 39.61 | 0.38 | 6.53 | 0 |
|  | 588 | 22.66 | 49.63 | 57.45 | 37.35 | 1.15 | 4.04 | 0 |
| **Average** |  | **20.75** | **47.09** | **60.17** | **35.24** | **0.57** | **4.01** | **0** |
| D1NAMO | 007 | 5.9 | 12.40 | 94.00 | 6.00 | 0 | 0 | 0 |
|  | 001 | 7.1 | 19.37 | 92.98 | 6.67 | 0 | 0.35 | 0 |
|  | 004 | 8.5 | 23.35 | 90.31 | 8.67 | 1.02 | 0 | 0 |
| **Average** |  | **7.17** | **18.37** | **92.43** | **7.11** | **0.34** | **0.12** | **0** |

**Table 5.** Results of ablation analysis on the OHIO dataset.

| Parameter | RMSE | MARD |
| --- | --- | --- |
| No HR | 27.73 | 9.70% |
| No cos_time_of_day | 16.65 | 7.96% |
| No sin_time_of_day | 15.75 | 7.42% |
| No insulin rate | 17.66 | 7.16% |
| No SDNN | 14.8932 | 6.80% |
| No RMSSD | 14.3083 | 6.64% |
| No Activity level | 15.10 | 6.39% |
| No pnn_50 | 14.1356 | 6.33% |
| No HRVTriIndex | 13.5246 | 6.17% |
| No calories | 13.92 | 6.10% |
| No meanRR | 13.5172 | 6.05% |
| No meanHR | 14.6809 | 6.05% |
| No BR | 13.58 | 6.04% |
| ALL | 12.41 | 5.90% |
| No SDHR | 13.6364 | 5.85% |
| No insulin slow | 12.39 | 5.83% |
| No nn_50 | 13.3251 | 5.71% |
| No TINN | 12.6997 | 5.55% |
| No insulin fast | 11.73 | 5.42% |

OHIO Dataset:

The average MARD across patients is 20.09%, reflecting higher prediction errors. The RMSE is 47.09, indicating greater deviations between predicted and actual glucose levels. Zone A accuracy averages 60.18%, with a significant portion of predictions in Zone B (35.24%) and minor but notable percentages in Zones C and D.

D1NAMO Dataset:

The average MARD is significantly lower at 7.17%, demonstrating superior prediction accuracy. The average RMSE is 18.38, showing reduced prediction deviations compared to the OHIO dataset. Zone A accuracy averages 92.43%, with most remaining predictions in Zone B (7.11%), negligible entries in Zones C and D, and none in Zone E.

All numbers except RMSE are expressed as a percentage.

The results show notable differences in accuracy and reliability between the OHIO and D1NAMO datasets. These differences are due to the type of biomarkers and the frequency of data collection. Unlike the lower-frequency OHIO dataset, the D1NAMO dataset collects high-frequency data over a short period, enabling detailed feature extraction. Additionally, OHIO collected heart rate (HR) data, while D1NAMO collected heart rate variability (HRV) data, further influencing the results.

### Ablation Analysis

The ablation analysis for Patient 007 in the D1NAMO dataset is shown in Table 3 as a heat map. This analysis shows how removing specific features affects the model’s performance.

This analysis for Patient 007 reveals that the most critical features are HR and Insulin Rate, as removing them results in the largest increase in errors. The time of Day (sin_time_of_day, cos_time_of_day) is also necessary, as excluding it significantly impacts predictions.

Moderately important features include Activity Level, Breathing Rate (BR), and Calories Expended. These features have a smaller but noticeable effect when removed. The least essential features include Insulin Fast, Insulin Slow, and HRV, which have minimal or no positive impact. Removing HRV even improves accuracy slightly. Insulin Fast and Insulin Slow alone do not make a difference in the model, as the Insulin Rate captures the combined effect.

The ablation analysis for the 30-minute forecasting model of Patient 570 from the OHIO dataset is presented in Table 6 as a heat map, with performance measured by MARD and RMSE.

**Table 6.** Ablation analysis for forecasting model on OHIO for Patient 570 for 30 minutes.

| Feature Removed | RMSE | MARD |
| --- | --- | --- |
| None | 6.644 | 2.29% |
| HR | 7.345 | 2.83% |
| Physical_activity | 6.673 | 2.46% |
| Operative_carbs | 6.326 | 2.14% |
| Active_insulin | 6.488 | 2.23% |
| Skin_temperature | 6.426 | 2.06% |
| Sleep_activity | 6.640 | 2.31% |
| Basis_steps | 6.549 | 2.26% |
| Sin_time_of_day | 6.353 | 2.06% |
| Cos_time_of_day | 6.848 | 2.31% |

The model has an RMSE of 6.644 and a MARD of 2.29%, with all features included. Removing specific features shows their importance to the model. Excluding Heart Rate (HR) causes a significant drop in performance, increasing RMSE to 7.345 and MARD to 2.83%, highlighting HR as a critical feature. Similarly, removing the Cosine of Time of Day (Cos_time_of_day) increases the RMSE to 6.848 and the MARD to 2.31%, underscoring the importance of time-related patterns.

Figure 4 compares Actual Glucose Levels (CGM data) and Predicted Glucose Levels (vBGM values) over time for the D1NAMO Patient 007. The MARD is 5.9%, RMSE is 12.40, and the Zone A accuracy is 94%. Both curves exhibit similar patterns, with the predicted glucose values closely following the actual glucose values across the time series, indicating strong alignment between actual and modeled glucose dynamics. The prediction model effectively captures the peaks and valleys in glucose levels, demonstrating its responsiveness to rapid changes. Slight deviations are visible, particularly at sharp transitions, where the predicted glucose levels lag slightly behind the actual values, suggesting room for improvement in handling extreme changes and demonstrating robustness across a wide glucose range (approximately 50–400 mg/dL).

**Figure 4.**
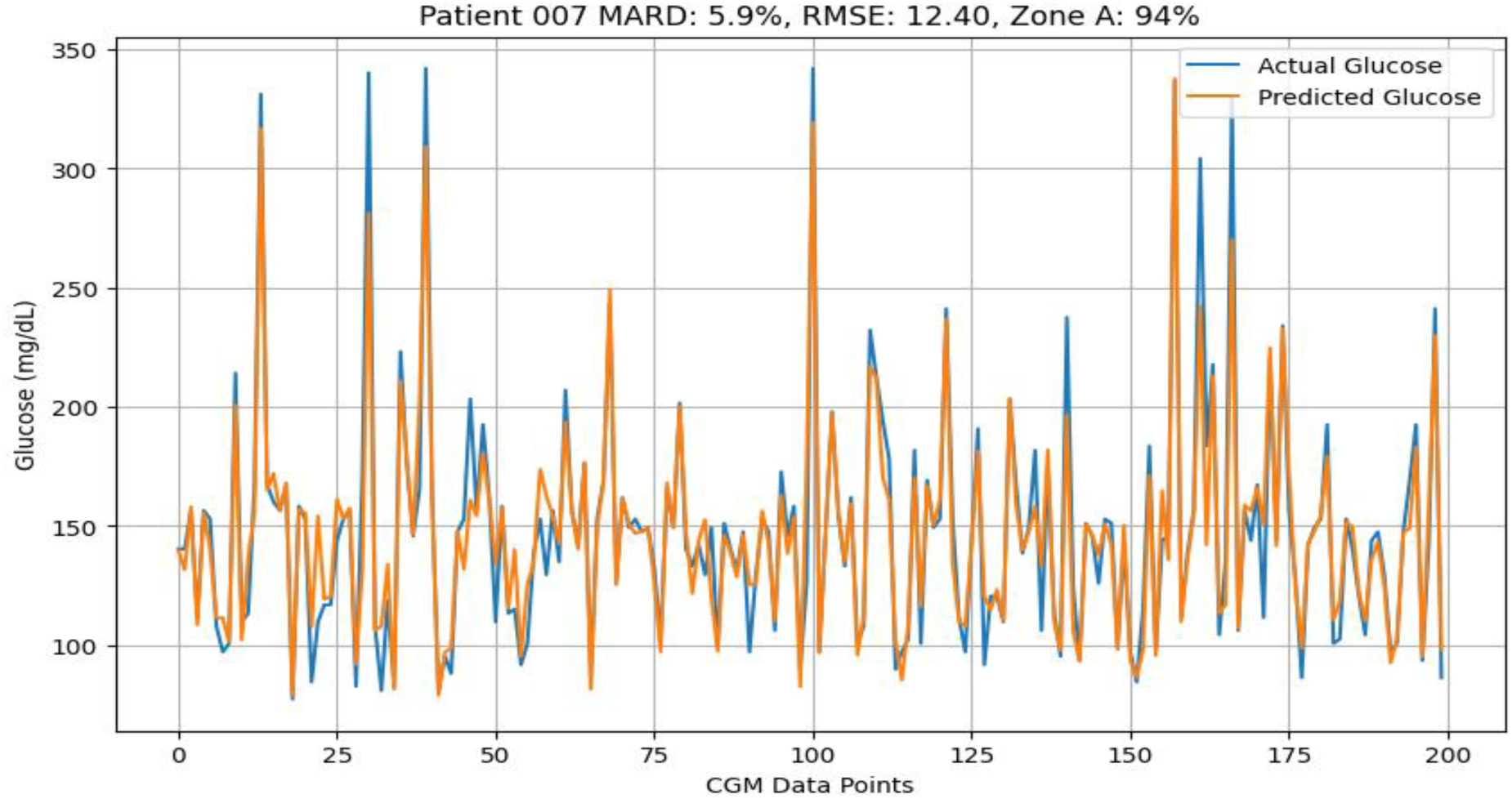
Actual CGM glucose values versus vBGM predicted values over time.

Figure 5 illustrates the Clarke Error Grid for Patient 007, specifically for vBGM predictions. The plot shows that most observations are in Zone A (94%), a few are in Zone B (6%), and none are in Zone C, D, or E. This distribution points to the model’s clinical accuracy.

**Figure 5.**
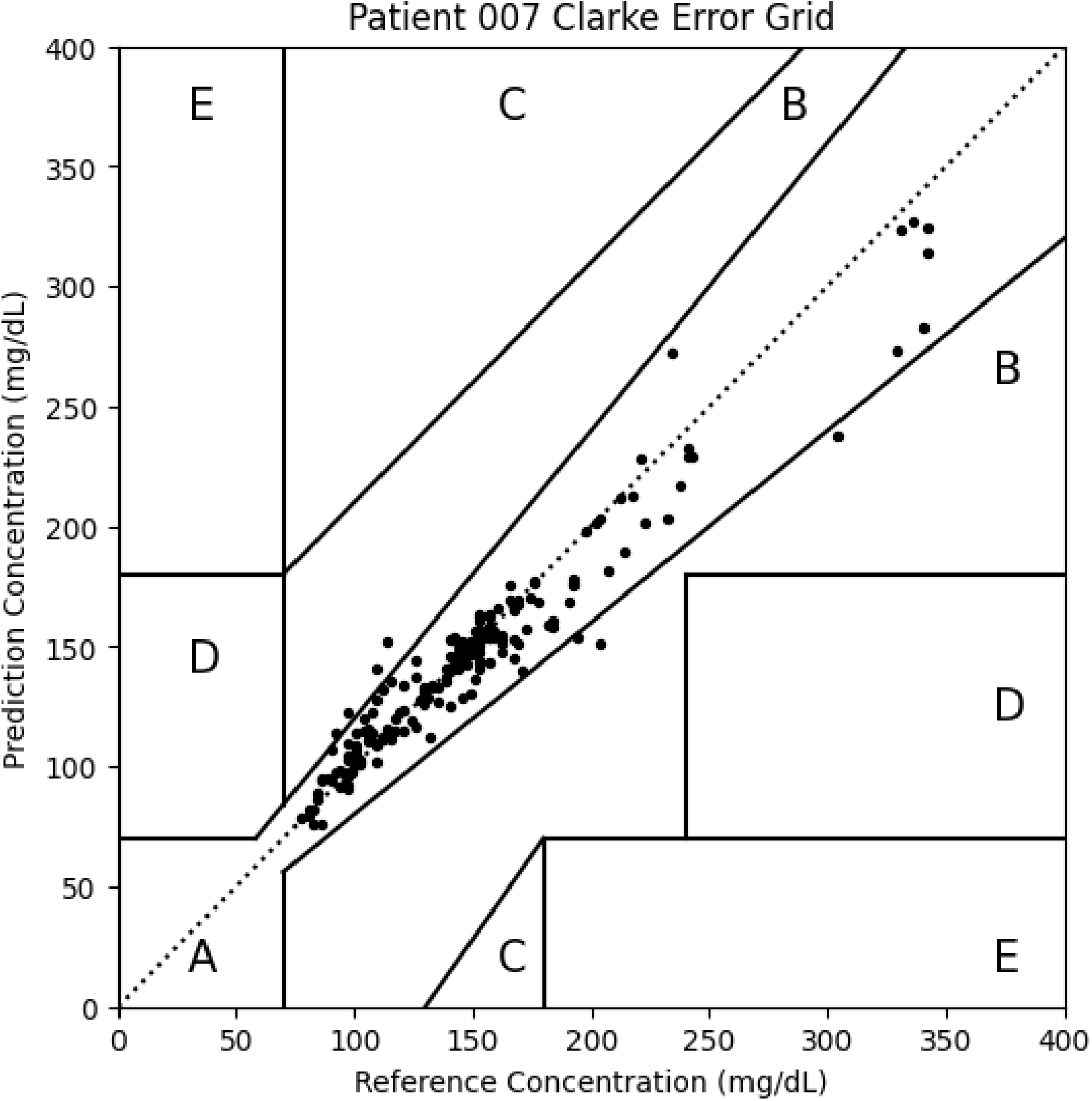
Clarke error grid for Patient 007 for vBGM predictions.

## Discussion and Conclusion

The CCA analysis of the D1NAMO dataset identifies key physiological metrics that contribute to glucose level predictions. Heart Rate, Insulin Rate, and temporal features emerge as significant predictors, particularly for specific patients, such as Patient 004 and Patient 007. Metrics with weaker or inconsistent correlations, such as RMSSD and R_Peaks, highlight the importance of carefully selecting features for predictive models. These findings highlight the significance of patient-specific variability and the targeted application of physiological metrics to enhance glucose prediction accuracy.

The Dual Temporal Recurrent Ensemble (DTRE) model demonstrates remarkable advancements in multi-horizon glucose forecasting, setting a new standard in personalized diabetes management. Across all forecasting horizons, the DTRE model consistently outperforms existing models on key metrics, including Mean Absolute Relative Difference (MARD), Root Mean Squared Error (RMSE), and Zone A Accuracy from the Clarke Error Grid. For instance, at the 30-minute horizon, the DTRE model achieves a MARD of 7.6% ± 3.4%, significantly surpassing the Zhu model’s 8.7%. Similarly, its RMSE is 16.49 ± 5.504, reflecting improved precision in capturing glucose dynamics.

In all aspects, the OHIO dataset outperformed the D1NAMO dataset for the forecasting model. The OHIO dataset had a longer data collection time and more data points. Therefore, the forecasting model will benefit from longer-term data collection to achieve more accurate forecasts.

On the high-resolution D1NAMO dataset, the DTRE model achieves a Zone A Accuracy of 92.43%, significantly outperforming the OHIO dataset’s 60.18%, highlighting the critical role of high-frequency data collection. The MARD for vBGM was lower than the Abbott Labs FreeStyle Libre 3 CGM’s MARD of 7.9% (Abbott, 2023). Anderson, Rivera, and Thompson (2024) argue that virtual glucose monitors leveraging advanced AI architectures can significantly improve accessibility and affordability for patients, especially in resource-limited settings.

The reverse ablation analysis for patient 007 showed that Heart Rate, Insulin Rate, and Time of Day are essential for accurate glucose predictions. Other features contribute less, and HRV may introduce additional noise for this patient. For the 30-minute forecasting model, Heart Rate and Cosine of Time of Day are the most essential features, as their removal significantly affects the model’s performance. Features like skin temperature, sleep activity, and basis steps have minimal impact and could be removed from this patient’s model. However, a more thorough analysis of all patients should be conducted before making this decision.

Despite its superior performance, opportunities for improvement remain. Incorporating the Glycemic Index into carbohydrate modeling and refining insulin equations to handle hybrid doses and brand-specific variations could enhance predictive accuracy. Expanding datasets to include diverse populations and automating data collection processes via APIs would improve model generalizability and reduce errors.

## Limitations

While the Dual Temporal Recurrent Ensemble (DTRE) model demonstrates significant advancements in glucose prediction, several limitations should be acknowledged to guide future research and practical implementation:

### Dataset Size and Diversity

While robust, the datasets used in this study are limited in size and demographic diversity. The OHIO and D1NAMO datasets primarily include data from a restricted number of patients, which may not fully capture the variability in glucose dynamics across broader populations. Ethnicity, age, gender, and regional dietary habits influence glucose regulation, but these factors are underrepresented in current datasets. Expanding the dataset to include more diverse populations will improve model generalizability and clinical applicability.

### Computational Requirements

The DTRE takes about 30 minutes to run on Kaggle. While this is not a significant burden, it may become an issue for many patients and necessitate periodic training of all models. This may pose challenges in settings with limited computational infrastructure. Optimizing the model to reduce complexity without sacrificing performance will be necessary for scalability.

### Edge Cases and Failure Modes

The model’s performance in edge cases, such as during extreme physical activity, illness, or erratic dietary habits, remains to be fully validated. These scenarios often introduce rapid glucose fluctuations that the current feature set or model architecture may not adequately capture. Identifying and addressing failure modes, such as inaccurate predictions during these edge cases, will enhance the model’s reliability in diverse conditions.

## Feature Engineering

Feature engineering played a pivotal role in developing the DTRE model, significantly enhancing its predictive capabilities. The innovative carbohydrate modeling, utilizing a piecewise function, helped capture the nonlinear impact of dietary intake on glucose levels. However, the current approach does not account for the Glycemic Index, a critical factor influencing the rate of glucose conversion. Future iterations should integrate this metric to improve predictions of postprandial glucose dynamics.

Insulin modeling was another cornerstone of the feature engineering process. While the model effectively incorporates insulin data, it does not yet fully account for hybrid insulin doses (a combination of fast-acting and slow-acting insulin) or brand-specific models. This enhancement will provide more accurate insulin-on-board (IOB) estimates, enabling better prediction of glucose fluctuations.

Physical activity was modeled using basic metrics, but there is room for refinement. Incorporating heart rate, heart points, and specific exercise types, such as aerobic or resistance training, could improve the model’s ability to predict activity-induced glucose changes. Modern fitness devices provide data on stress and sleep, which influence glucose dynamics, and their inclusion would further enhance the model’s predictive power.

Finally, automated data collection through Application Programming Interfaces (APIs) for food intake, physical activity, and other parameters could address current limitations in manual data entry. This improvement would reduce errors, minimize missing data, and ensure more accurate, timely data inputs, thereby increasing the model’s responsiveness and reliability.

## Future Work

Future developments of the DTRE model and vBGM should focus on the following areas to enhance their utility and performance:

1. Integration with Wearable Devices: Integrating the vBGM with wearable devices, such as smartwatches and fitness trackers, could improve the accuracy and timeliness of data collection. Wearables offer non-invasive measurements of heart rate variability (HRV), activity levels, and stress, thereby reducing user burden and increasing data frequency.
2. Real-Time Implementation Considerations: Real-time glucose prediction necessitates the use of optimized algorithms to process high-frequency data with minimal latency. Future work should explore edge computing and robust error handling to improve the model’s responsiveness in dynamic environments.
3. Potential for Personalization: Personalizing the model to account for individual glycemic responses, insulin sensitivity, and lifestyle factors could significantly enhance its clinical applicability. Tailored predictions would empower users with more accurate and actionable insights.
4. Additional Features to Explore: Incorporating novel biomarkers, such as stress levels, sleep quality, dietary diversity, and environmental factors (e.g., temperature and altitude), could improve the model’s predictive power and applicability across diverse scenarios.
5. Data collection time: Most modern CGMs, such as the Libre 3 from Abbott Labs, span 14 days. Data should be collected for at least 14 days and at high frequency to obtain the most accurate data possible for that patient.
6. Combine it with Fingerstick: Many patients may be unable to afford a CGM. This model should be enhanced with periodic fingerstick measurements to train and validate the model. The degree of accuracy would be less.
7. All this research is done for Type 1 diabetes. This model should be tested on type 2 diabetes patients with and without insulin dependency.

## Conclusion

The DTRE model establishes a new benchmark in multi-horizon glucose prediction, offering unparalleled accuracy and clinical reliability. Its performance metrics, such as a MARD of 7.6% ± 3.4% and Zone A Accuracy of 99.79% at 30 minutes, demonstrate its superiority over existing models and commercial continuous glucose monitoring (CGM) devices like FreeStyle Libre 3 (MARD: 7.9%) and Dexcom G7 (MARD: 8.2%).

The DTRE model bridges the gap between real-time monitoring and extended forecasting by leveraging advanced feature engineering and hybrid architectures. Its ability to track glucose dynamics and predict future trends empowers patients and healthcare providers to make proactive interventions, reducing the risks of hypoglycemia and hyperglycemia and enhancing safety.

Nonetheless, this study highlights areas for future research, including incorporating the Glycemic Index, hybrid insulin modeling, and stress and sleep metrics. Expanding datasets to include diverse populations and integrating modern wearable devices with high-resolution sensors will refine the model’s accuracy and applicability.

The DTRE model represents a transformative advancement in diabetes management. Its dual real-time and predictive monitoring capabilities, along with robust feature engineering, pave the way for accessible, precise glucose management tools. Future work aimed at integrating wearable technology, enabling real-time use, and enhancing personalization will ensure that this innovation continues to evolve, benefiting individuals globally.

## Acknowledgment

I am deeply grateful to the subjects who contributed their data to the OHIOT1DM and D1NAMO datasets, making this project possible. Their participation was vital to the success of this research. I also appreciate the researchers who have advanced our understanding of this critical disease.

Lastly, I am deeply grateful to my family for their constant support. Their encouragement inspired me to create a tool to help Type 1 diabetic patients manage their condition more effectively.

## Appendix

Figures for extended horizons (30, 60, 120 minutes) included below

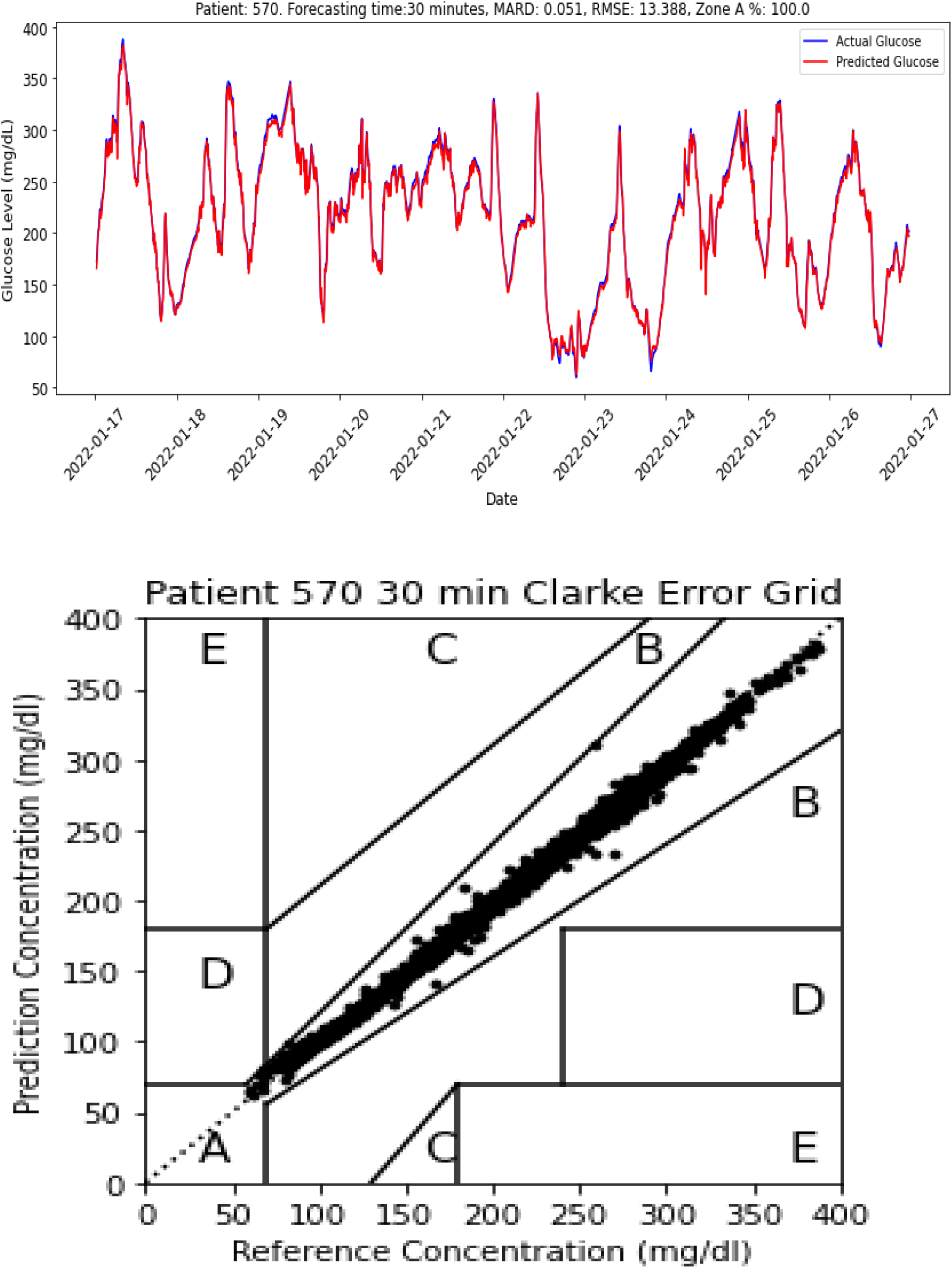

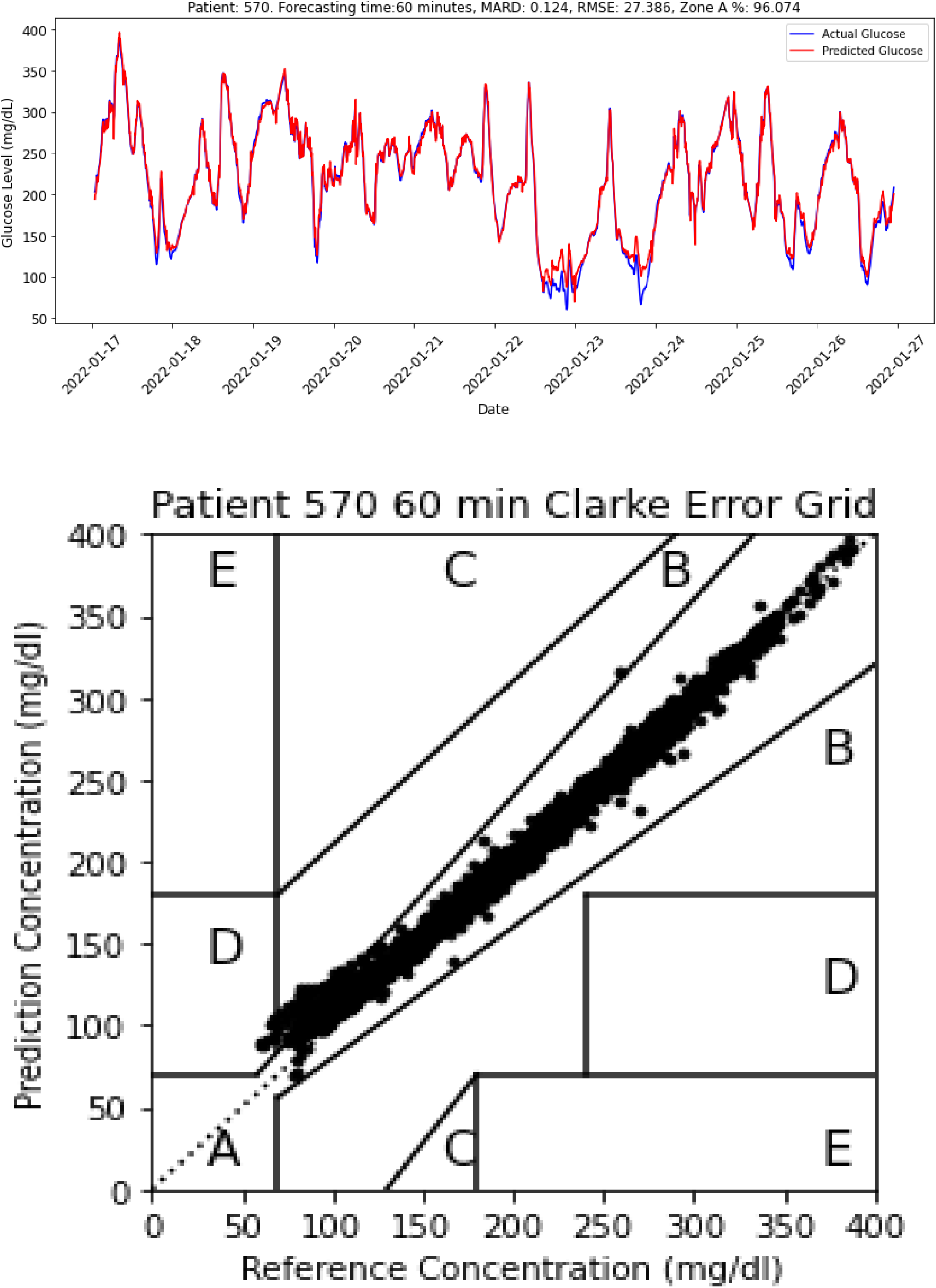

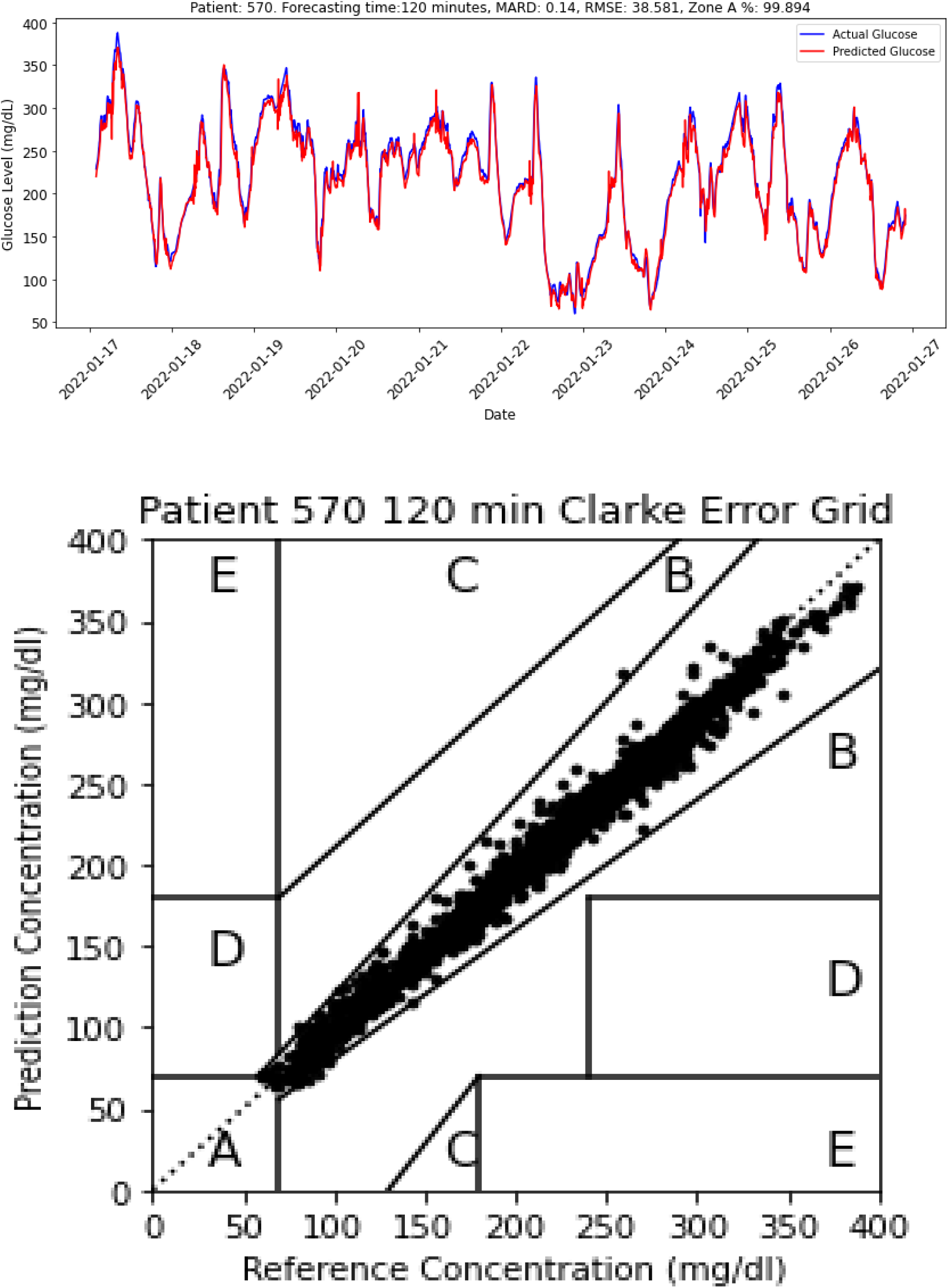

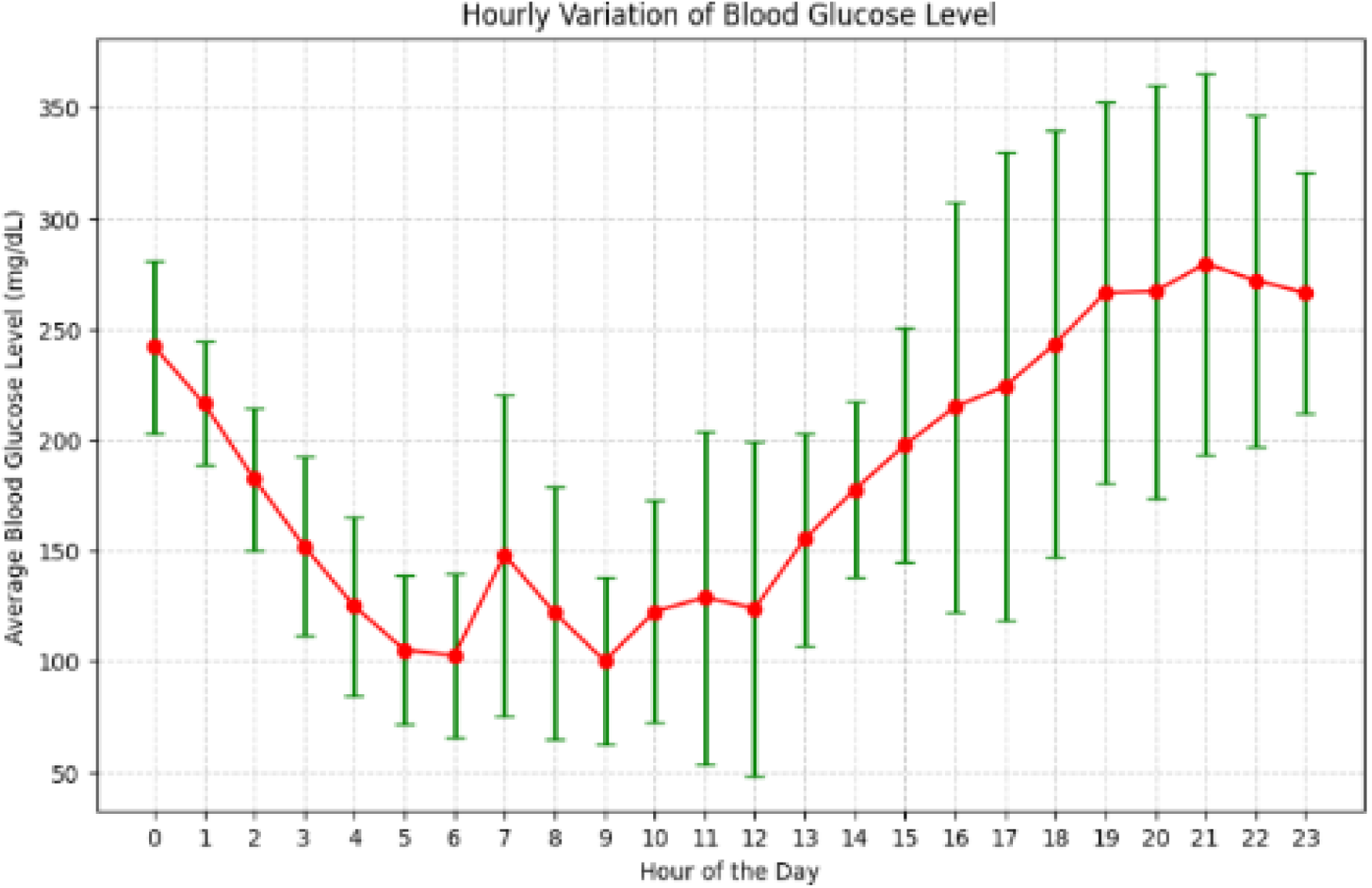

## References

Abbott. (2023). Welcome to the world of MARD: Transforming diabetes care with the FreeStyle Libre 3. Abbott. https://www.abbott.com/corpnewsroom/diabetes-care/welcome-to-the-MARD-world-of-diabetes-care.html

American Diabetes Association. (2024). Standards of medical care in diabetes—2024. Diabetes Care, 47(Supplement 1). 10.2337/dc24-S001

Anderson, B., Rivera, J., & Thompson, D. (2024). Beyond CGM: Exploring AI-driven virtual glucose monitors for affordable diabetes care. Journal of Medical Internet Research, 26, e45239. 10.2196/45239

Bhimireddy, A., Sinha, P., Oluwalade, B., Gichoya, J. W., & Purkayastha, S. (2020). Blood glucose level prediction as time-series modeling using sequence-to-sequence neural networks. CEUR Workshop Proceedings.

Butt, H., Khosa, I., & Iftikhar, M. A. (2023). Feature Transformation for Efficient Blood Glucose Prediction in Type 1 Diabetes Mellitus Patients. Diagnostics (Basel, Switzerland, 13(3), 340. 10.3390/diagnostics13030340

Clarke, W. L., Cox, D., Gonder-Frederick, L. A., Carter, W., & Pohl, S. L. (1987). Evaluating clinical accuracy of systems for self-monitoring of blood glucose. Diabetes Care, 10(5), 622–628. 10.2337/diacare.10.5.622

Dubosson, F., Ranvier, J.-E., Bromuri, S., Calbimonte, J.-P., Ruiz, J., & Schumacher, M. (2018). The open D1NAMO dataset: A multi-modal dataset for research on non-invasive type 1 diabetes management. Informatics in Medicine Unlocked, 13, 10–101. 10.1016/j.imu.2018.09.003

El-Khatib, F. H., Jiang, J., Damiano, E. R., & Russell, S. J. (2010). A computational model of the human glucose-insulin regulatory system: Applications in closed-loop control. Journal of Diabetes Science and Technology, 4(6), 1327–1338. 10.1177/193229681000400601

Freckmann, G., Schmid, C., Baumstark, A., Pleus, S., Link, M., & Haug, C. (2012). System accuracy evaluation of 43 blood glucose monitoring systems for self-monitoring of blood glucose according to ISO 15197. Diabetes Technology & Therapeutics, 14(2), 113–124. 10.1089/dia.2011.0153

Gomes, P., Margaritoff, P., & Silva, H. (2019). pyHRV: Development and evaluation of an open-source Python toolbox for heart rate variability (HRV). Proceedings of the International Conference on Electrical, Electronic and Computing Engineering (IcETRAN), 822–828. Retrieved from https://github.com/PGomes92/pyhrv

Guan Z., Li H., Liu R., Cai C., Liu Y., Li J., Wang X., Huang S., Wu L., Liu D., Yu S., Wang Z., Shu J., Hou X., Yang X., Jia W., Sheng B. (2023). Artificial intelligence in diabetes management: Advancements, opportunities, and challenges. Cell Rep Med, 4(10), 101213. 10.1016/j.xcrm.2023.101213

International Diabetes Federation. (2022). Diabetes Atlas.

Kim, H., Lee, J., & Choi, S. (2023). A comparative study of hybrid machine learning models for blood glucose prediction. Journal of Diabetes Science and Technology, 17(3), 112–121. 10.1177/19322968231103892

Kim, J., Choi, Y., & Oh, S. (2020). Prediction of blood glucose levels using recurrent neural networks. Diabetes Technology & Therapeutics, 22(10), 805–812. 10.1089/dia.2020.0035

Liu, J., Gao, Q., & Chen, H. (2024). Feature engineering techniques in AI-driven diabetes care: A review of the state-of-the-art. Biomedical Signal Processing and Control, 85, 105541. 10.1016/j.bspc.2023.105541

Marling, C., & Bunescu, R. (2020). The OHIOT1DM Dataset for Blood Glucose Level Prediction: Update 2020. CEUR Workshop Proceedings, 2675, 71–74.

Masterarbeit von Gizem Gülesir, Tag der Einreichung, Sebastian Kauschke, and Gizem Gülesir. (2018). Predicting blood glucose levels of diabetes patients.

Patel, R., Johnson, A., & Singh, M. (2023). Advancing non-invasive glucose monitoring: The role of heart rate variability in predicting glycemic fluctuations. Artificial Intelligence in Medicine, 136, 102438. 10.1016/j.artmed.2023.102438

Qingnan Sun, Marko Jankovic, Lia Bally, & Stavroula Mougiakakou. (2018). Predicting blood glucose with an LSTM and bi-LSTM-based deep neural network. Proceedings of IEEE EMB Annual Conference, 1–5.

Smith, A., Brown, K., & Wilson, P. (2024). Integrating physiological and behavioral data for personalized diabetes management: Opportunities and challenges. Diabetes Technology & Therapeutics, 26(1), 45–54. 10.1089/dia.2023.0369

Sun, Q., Jankovic, M., Bally, L., & Mougiakakou, S. (2018). Predicting blood glucose with an LSTM and bi-LSTM-based deep neural network. Proceedings of IEEE EMB Annual Conference, 1–5.

T. E. Idriss, A. Idri, I. Abnane, & Z. Bakkoury. (2019). Predicting blood glucose using an LSTM neural network. 2019 Federated Conference on Computer Science and Information Systems (FedCSIS), 35–41.

Willmott, C. J., & Matsuura, K. (2005). Advantages of the mean absolute error (MAE) over the root mean square error (RMSE) in assessing average model performance. Climate Research, 30(1), 79–82. 10.3354/cr030079

Zhu, T., et al. (2021). Machine learning approaches for predicting blood glucose levels. Journal of Diabetes Science and Technology, 15(3), 575–582.

